# Stopping Speed as State, Not Trait: Exploring Within-Animal Varying Stopping Speeds in a Multi-Session Stop-Signal Task

**DOI:** 10.1101/2024.09.05.611370

**Authors:** Jordi ter Horst, Michael X Cohen, Bernhard Englitz

## Abstract

Being able to reactively stop ongoing movements is important for safe navigation through the environment. Reactive stopping is typically studied using the stop-signal task, where participants are occasionally instructed to stop initiated movements. The speed of stopping, also referred to as the stop-signal reaction time (SSRT), is not observable because successful stopping lacks a response, but can be estimated. Researchers most often acquire one session of data per participant to estimate the speed of stopping, but sometimes more sessions of data are acquired to maximize the signal-to-noise ratio, for example when the task is combined with neural recordings such as electrophysiology. However, it is unknown whether the estimated stopping speed is a fixed trait or a state that can vary under identical experimental conditions. In this study, we investigate whether a separately estimated SSRT for each acquired session is statistically meaningful compared to estimating an across-session SSRT, by collecting many sessions in which male rats performed a stop-signal task. Results revealed that within-animal stopping speeds meaningfully changed from session to session and were not following a trend over time (e.g., due to task learning). Single-session SSRT estimates with lower reliabilities were associated with higher go trial response time variabilities, lower skewness levels of the go trial response time distribution, and lower stop accuracies. We also explored which factors explained changing SSRTs, and showed that motivation, shared motor dynamics, and attention could play a role. In conclusion, we encourage researchers to treat SSRTs as state-like variables when collecting multi-session stop-signal task data, as our results have convincingly shown that stopping speeds are far from trait-like under identical experimental conditions. This session-by-session approach will help future research in which neural signatures of reactive stopping need to be extracted in a time-precise manner, because time-locking stop-related neural activity to session-specific SSRTs is expected to capture the signature more precisely as opposed to an across-session SSRT.

## Introduction

### Reactive stopping in the real world

Being able to suddenly stop movements is a vital skill for navigating safely through the environment. Car drivers, bikers and other traffic users may suddenly cross your way while you are running around the block. Potential threatening external stimuli like these are taken very seriously by the brain; somehow it manages to reactively stop the ongoing running in a split second to prevent collision, without you even thinking about it. Reactive stopping, as defined here, is extremely fast and driven by external sensory input. The field of neuroscience has put many efforts into understanding how reactive stopping is implemented in the mammalian brain, but has not been conclusive about the specific neural mechanisms underlying the quick capability of stopping ongoing actions.

### Reactive stopping in the lab

Reactive stopping is typically studied with the stop-signal task, in which participants are presented with a short-lasting go-signal, requiring them to respond to this visual or auditory stimulus as quickly as possible by pressing a button (Logan & Cowan, 1984; Verbruggen & Logan, 2009; Verbruggen et al., 2019). Occasionally, the go-signal is followed by a stop-signal, instructing to cancel the ongoing motor plan of pressing the go-signal button (Figure 1). This task can be anywhere from easy to very difficult, depending on the delay between the go-signal and the stop-signal. This delay is called the stop-signal delay. Comparable to a traffic light turning red while you were about to cross the road, a shorter delay (i.e., earlier stop-signal presentation) makes it easier to still stop the ongoing plan or action, while a longer delay (i.e., later stop-signal presentation) makes it harder to stop in time. In an experimental lab setting, successfully stopping an ongoing action does not come with an observable button press because participants have to cancel the ongoing move towards the button. Therefore, researchers need to estimate the non-observable stopping time, also called stop-signal reaction time (SSRT). This is done by making use of observable information such as the time needed to press when only a go-signal is presented, and the probability of (erroneously) responding with a button press when a stop-signal was presented (Verbruggen et al., 2019).

**Figure 1.**
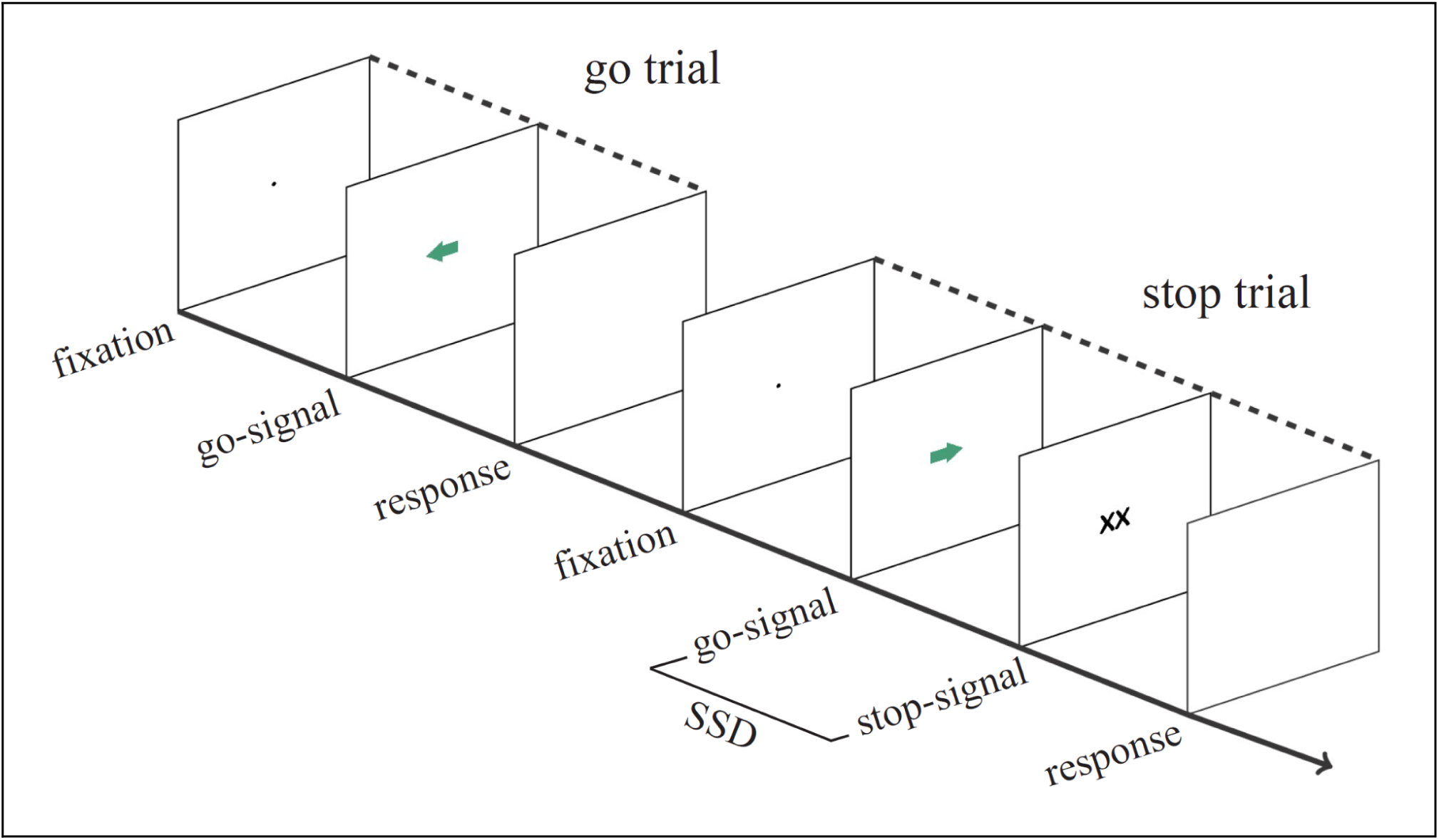
Typical stop-signal task for humans. After shortly fixating at a central position on a screen, a go-signal is presented instructing the participant to respond at the corresponding arrow direction side with a button press (go trial). Occasionally, this go-signal is followed by a stop-signal after a variable stop-signal delay (SSD), requiring the participant to not respond to the initially presented go-signal (stop trial). Taken and adapted from Figure 1 in Verbruggen et al. (2019).

### Is stopping speed a state or trait?

Quite often, researchers ask participants to perform the stop-signal task once and they need many participants for their study to acquire enough statistical power, as researchers are for example interested in whether a clinical group is slower at stopping than a non-clinical group. While this approach may be well-suited for the goal of the study, the downside is that it assumes that stopping speed is a trait that does not change from time to time. While cognitive development in childhood and adolescence is known to be associated with increased performance on stop-signal tasks, resulting in faster stopping speeds (Williams et al., 1999; Curley et al., 2018; Madsen et al., 2020), less is known about whether stopping speed is a fixed characteristic or a variable state after full cognitive development in non-clinical circumstances. Thunberg et al. (2024) have recently shown that test-retest reliability of SSRT estimates are low, even though SSRT estimates had high reliability within a session as demonstrated with high split-half reliability. However, to our knowledge there is no report out there that rigorously investigated how (in)variable stopping speeds are under identical experimental conditions in cognitively developed participants, and which factors drive potential instabilities in stopping speeds.

### Animal models to study stopping

As opposed to comparing stopping speeds of a clinical group with a non-clinical group, researchers often have to turn to animal models when they are interested in the neural mechanisms of stopping, especially when high spatial and temporal resolution are required. In these cases, researchers do not require many animals because within-subject, multi-session experimental designs are easier to perform with animals while maintaining enough statistical power. As a consequence, while many sessions are acquired, it remains unclear how one should handle multiple within-subject SSRT estimates. Are they statistically meaningful, reliable and useful?

### Our experiment and key findings

To this end, we trained male rats (N = 6) on a rodent version of the stop-signal task and investigated whether within-animal single-session SSRT estimates were statistically meaningful as compared to just estimating an SSRT across all acquired sessions as if stopping speed were a fixed trait of the animal. Overall, we show that stopping speed is not a fixed trait, but a state that changes from time to time. Within-animal single-session SSRTs varied substantially over the course of many sessions, for some animals more than others. As compared to the within-animal, across-session SSRTs, single-session SSRTs were much less reliable. In addition, we explain which factors may play a role in different degrees of single-session SSRT reliability, and which cognitive and neural mechanisms may underlie changing stopping speeds.

## Materials and Methods

### Animals

Six wild-type Long-Evans rats participated in this study, aged 9 weeks and weighing 250-320 grams at the start of behavioral training (Charles River Laboratories, Calco, Italy). Rats were housed pairwise in Makrolon type III cages (UNO B.V., Zevenaar, The Netherlands) with a reversed 12-hour day-night cycle in a temperature- and humidity-controlled room (21 ± 2°C, 60 ± 15%). As soon as the rats weighed more than 350 grams, they were housed in Makrolon type IVS cages to provide more horizontal space. As soon as the rats were implanted with electrodes (for another study, see chapter 3) they were housed individually, and the low conventional cage lid was replaced by a high cage lid to prevent damage to the implant. Corn cob granules were used as cage bedding, and sizzle bedding and a cardboard shelter were provided as cage enrichment. The rats were put on a restricted water intake schedule as soon as they acclimatized in the research facility. Every Monday to Friday morning the rats could get water in the behavioral task (∼5-8 mL, depending on performance), and in the afternoon they could drink *ad libitum* water for 30 minutes from a bottle. During weekend days, the rats received 30 grams of hydrogel (ClearH2O Inc., Westbrook, Maine, USA) each day, to keep the daily intake of water as stable as possible. Food pellets were provided *ad libitum* at all times. Weight and health were monitored on a daily basis. All animal procedures were approved by the Animal Welfare Body of the Radboud University Nijmegen and the Animal Experiment Committee (CCD No. AVD10300 2016 482, Project No. 2015-0129), according to national and international laws, to protect welfare under experimental conditions.

### Skinner box

After acclimation of two weeks in the research facility the rats started with the restricted water intake schedule and behavioral training. Training and testing took place in a custom-built Skinner box (inside dimensions: 25 × 27 × 25 cm), with one wall containing three nose-poke ports (bottom-left, bottom-center, bottom-right, see Figure 2). Each port had an infrared emitter and phototransistor (type L-53F3C and L-53P3C, peak 940 nm, Farnell B.V., Utrecht, The Netherlands) enabling continuous automatic detection of a nose-poke by the rat. In addition, the left and right port also contained green light-emitting diodes (type L-53SGD-5V, peak 565 nm, Farnell B.V., Utrecht, The Netherlands) for presenting visual stimuli, as well as a small silicone tube at each bottom of the port for providing 50 µL water drop rewards driven by solenoid pumps (The Lee Company, Westbrook, Connecticut, USA). Electronics needed for the task in the Skinner box were controlled by a computer with custom-written code in MATLAB (R2018b, The MathWorks Inc., Natick, MA, USA).

**Figure 2.**
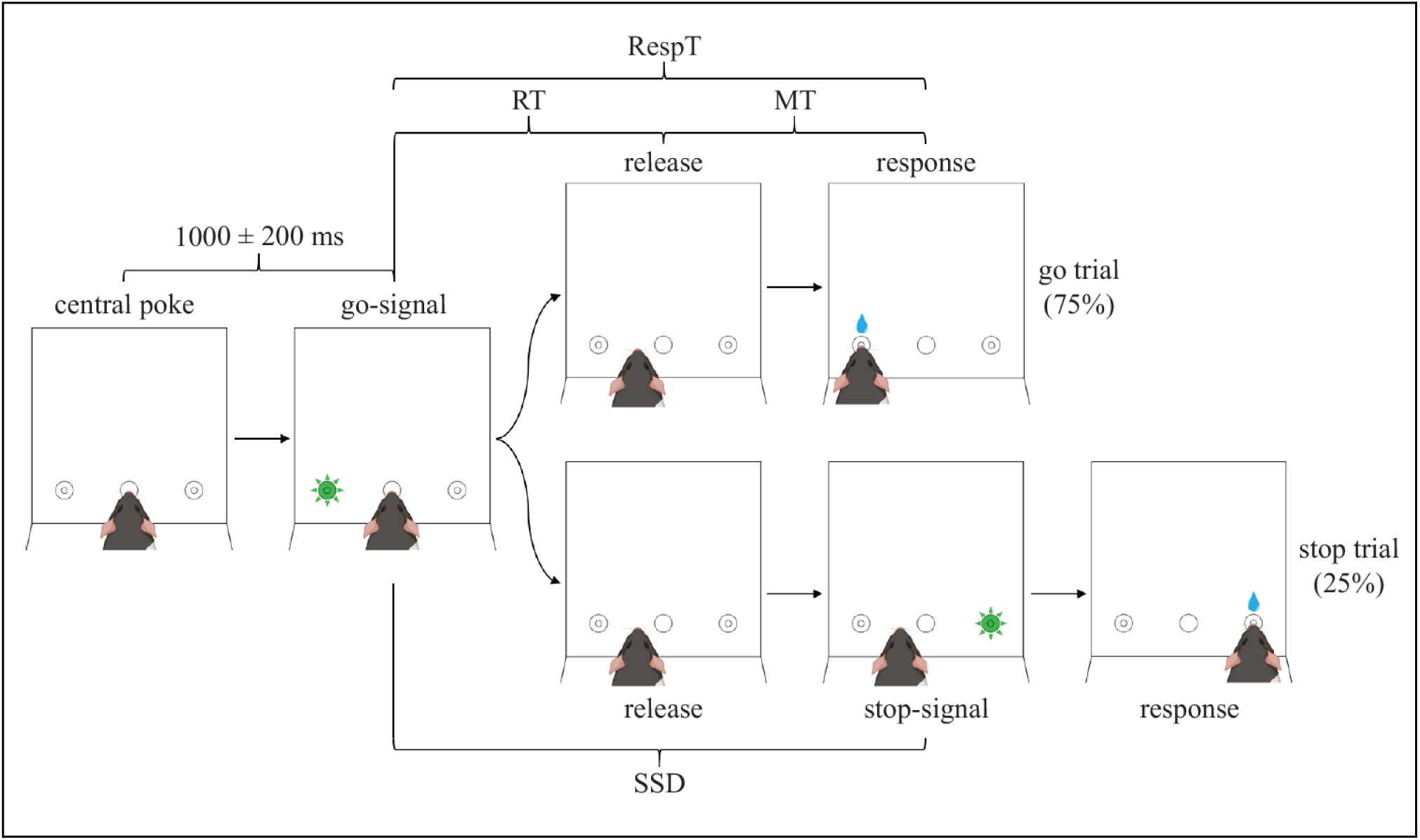
Optimized rodent version of stop-signal task in a Skinner box. Rats initiated trials by poking in a central nose-poke port for 1000 ± 200 ms, whereafter they were presented with a short visual go-signal on either the left or right side. In 75% of trials, no other signal followed and the rat had to respond at the go-signal side for a water drop reward. However, in 25% of the trials, the go-signal was followed by a visual stop-signal after a variable delay (stop-signal delay, SSD), which instructed the rat to stop moving to the go-signal side and respond at the stop-signal side to obtain a water drop reward. Only correct responses were rewarded during data acquisition. RT = release time; MT = movement time; RespT = response time.

### Training procedure stop-signal task

The rodent version of the stop-signal task was based on the behavioral tasks used by Feola et al. (2000) and Bryden et al. (2012), with optimizations taken from Verbruggen et al. (2019). The training procedure consisted of three main steps, namely 1) central nose-poke initiation, 2) unilateral cue discrimination, and 3) stop trial introduction. Rats were trained for maximally one hour or 200 trials each day, while having rest during weekend days. During the central nose-poke initiation phase, the rats had to learn to initiate a trial by poking their nose in the central port. On the first day of training, poking at the central port for the shortest time detectable was already enough to initiate a trial, causing the presentation of a light cue (go-signal) for 100 ms either on the left or right side with an immediate water drop reward provided at the corresponding side. As the reward was provided immediately after trial initiation the rat was not required to make a correct response yet. Despite not needing a correct response yet, trials were never terminated before a response was recorded, causing the rats to learn that they always had to respond in either the left or right port before they could initiate a new trial. This response requirement was kept at all training phases and data acquisition. The idea of this phase was to let the rat learn to associate central nose-poking with a positive outcome. Each day, the time required to stay in the central port (i.e., trial initiation time) was increased by 100 ms if the rat managed to initiate at least 100 trials in the previous session. When rats would release their nose from the central port before the required trial initiation time, the trial was not initiated and had to be re-initiated completely. This means premature responses as described in Verbruggen et al. (2019) are not possible. As soon as rats could initiate at least 100 trials in a session with 1000 ms initiation time, they proceeded to the second phase of training.

In the second phase, the rats had to learn to respond correctly to the go-signal before getting a reward. In practice this means the rat initiated a trial, received a go-signal at either the left or right side, and would only get a reward when the rat poked his nose in the port where the light was presented. This allowed the rats to learn to associate the go-signal side with the reward side. When the response was correct, the reward was immediately provided at the response port. No reward was provided after responding incorrectly. During this phase of training, the minimal time between the response at the lateral port and initiating a new trial (inter-trial interval) was gradually increased from 0 to 3 s in steps of 500 ms each session until 3 s was reached, to allow for proper separation of trials and to prevent rushing. To prevent go-signal anticipation, a jitter was slowly added to the trial initiation time from 0 to ±200 ms in steps of 50 ms every following session until a jitter of ±200 ms was reached. For each trial, the jitter value was randomly selected from a uniform distribution of numbers ranging from -200 to 200 ms with steps of 10 ms. As soon as the rats reached response accuracy above 80% for at least five consecutive days, they moved to the third and final phase of training.

In the final phase, stop trials were slowly introduced in addition to the go-only trials. On the first and second session of this final training phase the rats received 10% stop trials, the next three sessions 20% stop trials, and from the sixth day onwards 25% stop trials. In stop trials, the go-signal was followed by a stop-signal after a variable stop-signal delay (SSD). The stop-signal was given by a light on the other side than the go-signal side, and stayed illuminated until the rat made a response at either the left or right port (Figure 2). The SSD was determined separately for left and right stop trials through a staircase procedure, where it increased with 50 ms in case of a correct response, and decreased with 50 ms in case of an erroneous response. The SSD was not separately determined for two animals, and for two other animals only after ∼75% of their sessions were collected. This led us to exclude those four animals for this study, as we wanted consistency in the way the SSD was titrated for the particular purpose of this study. The starting SSD for stop trials was determined by averaging the SSDs from all stop trials from the previous session, for left and right stop trials separately. This staircase procedure is standard in human studies, ensures the collection of a wide range of SSDs and helps with obtaining a reliable stop-signal reaction time (SSRT) estimate (Verbruggen et al., 2019). Having a separately determined SSD for left and right stop trials ensures that the difficulty of a stop trial is always comparable between left and right stop trials, as a possible response bias may lead to an imbalanced accuracy for left versus right stop trials when there is a shared SSD. Go-signal side and stop trial occurrence were randomized and balanced as such that in each set of eight trials three left go trials, three right go trials, one left stop trial, and one right stop trial were randomly shuffled. Response bias was continuously checked by computing the percentage of left and right responses in the last 20 trials. When one side fell below 35%, the next trial was replaced by a go trial to that side to discourage response bias. As soon as the accuracy on stop trials floated around 50% and go trial accuracy was above 80% for five consecutive days, the rats were ready for electrode implantation. Training took approximately 6-8 weeks (30-40 sessions), and all rats achieved these criteria and were included in the subsequent recordings.

### Data acquisition

During the workweek rats performed the stop-signal task daily for one hour or maximally 200 trials, while intracranial local field potentials were recorded from the OFC and STN. This electrophysiological data is part of chapter 3 and is therefore not further discussed here. Parameters such as trial initiation time, release time, movement time, and stop-signal delay (in case of a stop trial) were automatically saved to disk on the computer controlling the Skinner box. Only data acquired after electrode implantation was used for behavioral analyses, so training data is not incorporated.

### Behavioral data preparation

Analyses of behavioral data were done with custom-written code in MATLAB software (R2018b, The MathWorks Inc., Natick, MA, USA). For each trial, release time was defined as the time between go-signal onset and release from the central port, movement time was defined as the time between release from the central port and response at either the left or right port, and summed together they form response time. Importantly, response time is the homologue of reaction time in human stop-signal tasks, as in human tasks reaction time is defined as the time between go-signal and response. In addition, for stop trials the stop-signal delay (SSD) was defined as the delay between go-signal onset and stop-signal onset (Figure 2). To deal with anticipatory releases (although discouraged with a jitter for go-signal onset), a local minimum in the smoothed bimodal release time distribution was identified and used as the lower bound for trial removal, as this portion of releases was anticipatory instead of reactive and attentive to the go-signal. The maximal lower bound was set at 100 ms, and the upper bound was set at three standard deviations above the mean release time. As no time limit was set for movement times, trials with movement times above five seconds were removed initially, whereafter the upper bound was set at three standard deviations above the mean for the remaining trials. Across the six included animals, trial removal based on release times and movement times resulted in an average removal of 15.1% of trials (95% CI [12.6 17.6]). The remaining trials were used for computing average release times, movement times, response times, stop-signal delays and stop-signal reaction times per session.

### Session exclusion criteria

Sessions were excluded based on five different exclusion criteria. Sessions were excluded when: 1) the average response time on unsuccessful stop trials was larger than the average response time on go trials, as those sessions violated the independence assumption from the horse-race model (Verbruggen et al., 2019) — 3.3% (95% CI [1.0 5.6]); 2) the response accuracy on stop trials was lower than 25% or higher than 75%, for left and right stop trials separately — 8.9% (95% CI [5.0 12.8]); 3) there were less than 10 stop trials presented on each side — 10.2% (95% CI [4.9 15.5]); 4) the response accuracy on go trials is higher than the accuracy on stop trials, left and right trials separately — 6.2% (95% CI [4.2 8.2]); 5) the left or right SSRT estimate of that session was lower than 100 ms — 1.2% (95% CI [-0.3 2.7]). Across animals, these criteria led to removal of 22.5% of sessions (95% CI [17.6 27.4]).

### SSRT estimation

Following the principles of the independent horse-race model, the unobservable stop-signal reaction time was estimated with the integration method (Verbruggen et al., 2019) for each session separately. In short, response times on go trials were sorted from shortest to longest, and the n-th response time matching the probability of failed stopping, —p(failed stopping)—, was found by multiplying p(failed stopping) from that session with the number of go trials in that session (see Figure 3 for a visual representation of go trial response times and how they relate to p(failed stopping), SSD and SSRT). Next, the SSRT was computed by subtracting the average SSD of that session from the n-th response time (nthRespTgo). As the SSD was separately determined for left and right stop trials, the SSRT was also separately estimated by using left and right response time distributions and SSDs. The response time distribution did not contain replacements for go omissions (go trials without a response), as responses were required on all trials. In addition, premature go responses (in our task releases before go-signal onset) are not part of the response time distribution either, as trials are not initialized in those cases (see training procedure). Both correctly and incorrectly performed go trials were used for the distribution. The across-session SSRT was estimated by putting data from all sessions together as if it was one big session, and then handled like in the above-mentioned procedure. Confidence intervals for SSRT estimates were computed with bootstrapping, by selecting the n-th response time matching p(failed stopping) from that session from a randomly sampled but sorted response time distribution retrieved from go trials. This n-th response time was in turn used to compute the associated SSRT by subtracting the SSD of that session from the n-th response time. After 1000 bootstraps, the 2.5% and 97.5% percentile of this set of SSRTs were extracted for the lower- and upperbound of the 95% confidence interval. The same bootstrapping procedure is used for the 95% confidence intervals of the across-session SSRT estimates.

**Figure 3.**
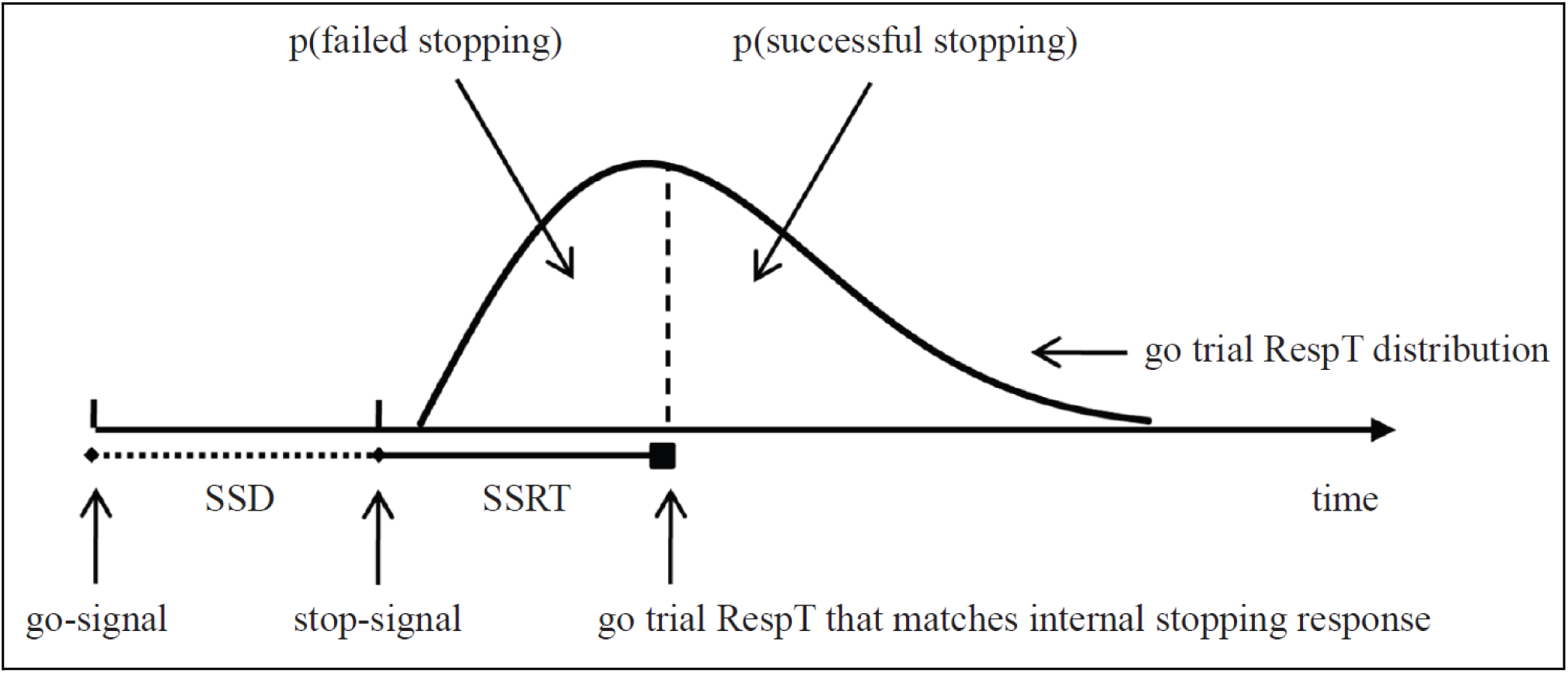
The independent horse-race model. The time it takes to stop (stop-signal reaction time; SSRT) can be estimated based on the go trial response time distribution, the SSD, and the probability of failed stopping. SSD = stop-signal delay; SSRT = stop-signal reaction time; RespT = response time. Importantly, RespT in our task is the homolog of reaction time in human stop-signal tasks, as in human tasks reaction time is defined as the time between go-signal and response. Taken and adapted from Figure 2 in Verbruggen and Logan (2009).

### Trend analysis of SSRT estimates

As SSRT estimations on a single-session basis could imply trends over time, we utilized a Matlab function called *RobustDetrend* (version 9.4.0) to get a general idea of potential SSRT development during the course of multiple sessions, for each animal and stop-signal side separately. This function is able to derive the best polynomial fit in a series of data while preserving peak features (Schivre, 2024). The polynomial order limit was set at 10, but only degree 0, 1 and 2 polynomials were found. We intentionally wanted to allow for peak features because SSRT estimations from sessions with smaller sample sizes may inherently have the tendency to show more legitimate outlier-like features, and we did not want to over-fit the data.

### Statistical meaningfulness of single-session SSRT estimates

To address whether single-session SSRTs (or state SSRTs) were statistically meaningful or just numerically different from across-session SSRT estimates (or trait SSRTs) due to noise and limited sample size, we computed the proportion of sessions where the single-session 95% confidence interval was overlapping with the across-session SSRT. Next, we investigated the degree of (dis)similarity between single-session SSRTs by computing the proportion of single-session SSRTs that were within the 95% confidence interval boundaries of other single-session SSRTs. As such, each session received a score between 0 and 100% and was used to compute an average across all single sessions. As a final step, the last analysis was repeated, but for neighboring sessions only, as this quantifies the speed at which proximate sessions can have significantly different SSRT estimates. For this, each session received a score depending on whether the single-session SSRT was not within neighboring 95% confidence intervals (score = 0%), within only one of them (score = 50%) or within both (score = 100%). In case of only one neighboring session (the first and last session), a score of either 0% or 100% was given (see Figure 4 for a visual representation of the three different analyses). All single-session percentages were averaged to obtain the percentage as illustrated in Figure 7.

**Figure 4.**
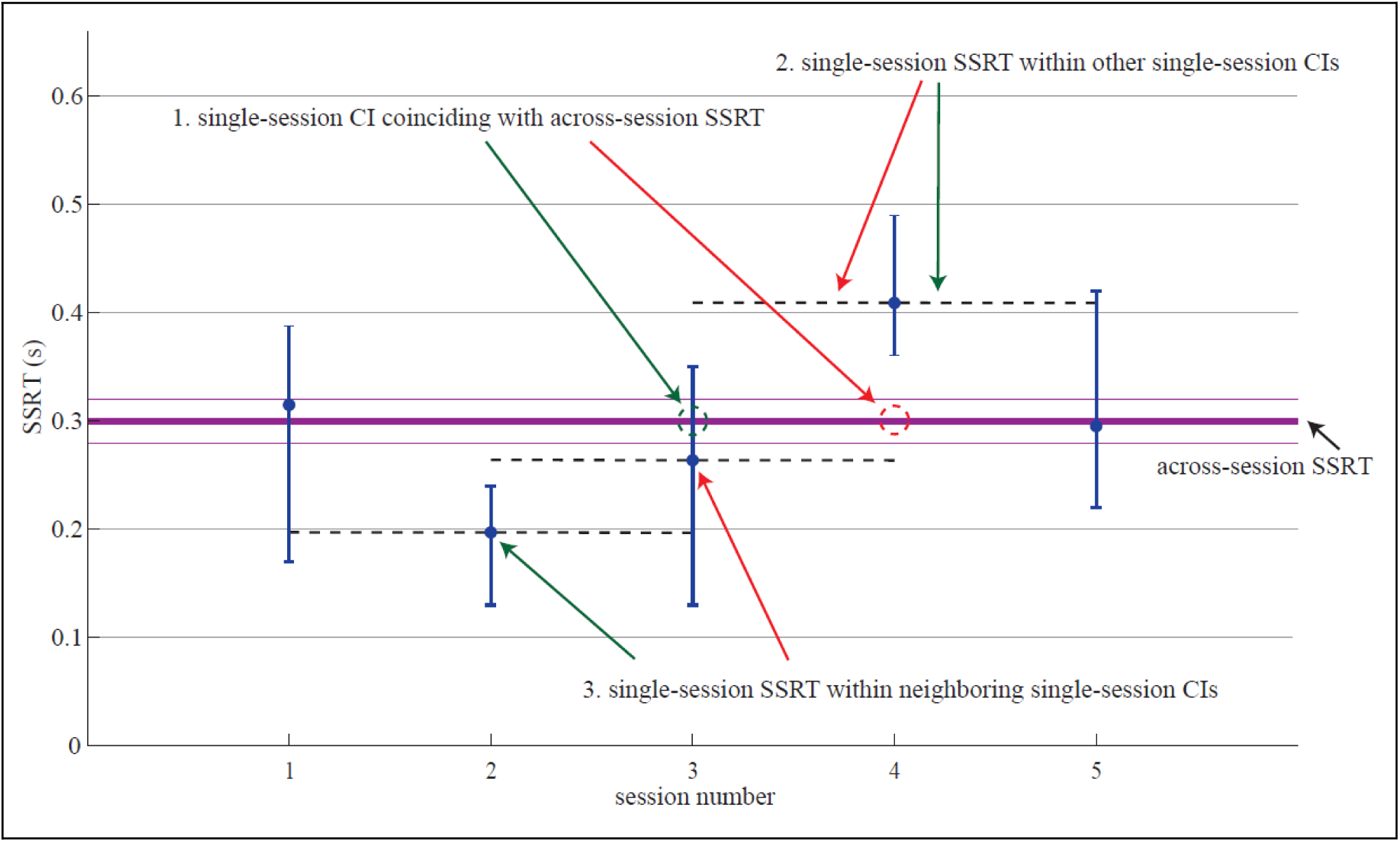
Fictive data to explain the different measures of statistical meaningfulness of single-session SSRT estimates. For each individual session, we checked whether 1) the single-session SSRT confidence interval (CI) was overlapping with the across-session SSRT estimate; 2) the single-session SSRT was within other single-session CIs; 3) the single-session SSRT was within neighboring single-session CIs. The purple thick line indicates the across-session SSRT with 95% confidence intervals indicated with thinner purple lines. Blue dots indicate single-session SSRT estimates with their corresponding 95% confidence intervals. Green arrows point to examples where the criteria are met as indicated, while red arrows point to examples where this is not the case. In this fictive example, criterion 1 would get a score of 60%, as three out of five sessions’ confidence intervals were overlapping with the across-session SSRT (session 1, 3 and 5). Criterion 2 would reach a score of 45%, as four sessions had an SSRT positioned within two other confidence intervals (session 1, 2, 3 and 5) while one session (session 4) had an SSRT positioned within one of the other four confidence intervals (4 x 50%, 1x 25% → 45%). Criterion 3 would receive a score of 50%, because session 1 and 2 were within their neighboring confidence intervals (score of 100%), session 3 and 5 were not within neighboring confidence intervals (score of 0%), and session 4 was only within one neighboring interval (score of 50%), leading to an average of 50% (2 x 100%, 2 x 0%, 1 x 50% → 50%).

These three different proportion calculations were repeated with detrended SSRT data to analyze if and how much proportions were affected by trends in SSRTs. We refrained from statistically testing whether detrending SSRTs affected these proportions because tests were heavily underpowered due to low sample size (N = 6). We qualitatively compared the proportions derived from unfiltered and detrended SSRTs.

### Reliability of single-session SSRTs

How certain one can be about a single-session SSRT estimate is quantified by the 95% confidence interval. Although it is inevitable to have larger confidence intervals as compared to the confidence obtained with across-session SSRTs, it is still worth investigating which factors were associated with smaller windows of confidence and hence more reliable single-session SSRT estimates. For each animal and trial side (left/right) we fitted a separate linear regression model with session-specific, independent variables: variability of go trial response times, skewness of the go trial response time distribution, stop-signal delay stabilization, stop accuracy and number of stop trials, and the 95% confidence interval width of SSRT as dependent variable. The model was fitted with the built-in Matlab function *fitlm* and included the intercept and main effect terms. As ordinary least squares models are highly sensitive to outliers, the model performed robust regression using the bisquare weighting function. This made the model less susceptible to disproportionate leverage from outliers. Variability of go trial response times was defined as the root mean square of demeaned go trial response times. Skewness of the go trial response time distribution was found with the built-in Matlab function *skewness*. Stop-signal delay stabilization was defined as the root mean square of demeaned stop-signal delays in the second half of the session. Stop accuracy was defined as the percentage of correct responses on stop trials, and lastly, the number of stop trials as the amount of stop trials that were originally in the uncleaned dataset. All variables were normalized to z-scores, so we could obtain standardized beta coefficients that were directly comparable.

To be able to visualize the relationship between each individual independent variable with the SSRT 95% CI width, we extracted the intercept and slope from simplified robust linear regression fits as described before, but with only one independent variable in the model. As such, we could include a linear line for each pair of independent variable and dependent variable in Figure 8A, for each animal separately. Lines were only solid when the independent variable in the full robust linear regression model had a p-value less than .05, and was dashed in case of non-significance. Standardized beta coefficients from all five independent variables and all six animals were represented with colors and summarized in heatmaps for left and right trials separately. A two-sided Wilcoxon signed-rank test was performed for each independent variable separately, and tested whether the standardized beta coefficients were significantly different from 0.

### SSD and nthRespTgo contribution to SSRT estimates

Because the non-observable SSRT was computed by subtracting the SSD from nthRespTgo, similar SSRTs could be obtained with many different combinations of SSD and nthRespTgo. Likewise, different SSRTs could be obtained with a constant SSD or nthRespTgo, while the other variable changes from session to session. To better understand the relationship between SSD, nthRespTgo and SSRT, we visualized these three variables together in Figure 9. We plotted the observable variables on the x- and y-axis, while using a diagonal for the across-session SSRT to illustrate how different sessions (and their corresponding SSD and nthRespTgo) relate to each other and the across-session SSRT. The average difference between single-session SSRTs and the across-session SSRT was computed for each animal and trial side (left/right) as a proxy for how spread out the single-session SSRT estimates were.

To statistically quantify the previously mentioned relationship, and to find out how much the SSD and nthRespTgo each contributed to session-to-session SSRT variance, we fitted a linear regression model for each animal and trial side (left/right) separately with SSRT as dependent variable and SSD and nthRespTgo as independent variables. The model was fitted with the built-in Matlab function *fitlm,* included the intercept and main effect terms, and made use of the bisquare weighting function to make the model more robust to outlier sessions. Standardized beta coefficients were extracted by z-scoring all input variables, so we could directly compare how much each variable’s variance contributed to session-to-session SSRT changes. A paired-samples t-test was used to statistically test whether there was a difference in how much each of two independent variables affected the SSRT. To this end, the magnitudes of standardized beta coefficients (i.e., ignoring the sign of the coefficients) from all animals’ robust linear regression models were used as paired samples.

### Possible cognitive and neural mechanisms driving session-to-session SSRT variability

Next to the obvious numerical predictors for session-to-session SSRT changes, SSD and nthRespTgo, we thought about which cognitive and neural mechanisms could possibly cause animals to have changing stopping speeds from session to session. Because the stop-signal task was optimized for estimating the SSRT and extracting neural stopping signatures, and not for investigating which factors contribute to within-animal SSRT variability across sessions, we approximated three different cognitive and neural mechanisms for changing SSRTs with the data that were available from the task: 1) motivation for quick rewards; 2) shared motor dynamics; 3) attention. Other possible contributing factors will be discussed theoretically in the discussion.

When animals are motivated for a quick reward, one would expect that they release fast after seeing the go-signal. Therefore, motivation for quick rewards was approximated with the release time on go trials (RTgo), and tested with linear regression by means of the built-in Matlab function *fitlm,* with a bisquare weighting function. Since this motivation could be expected at any stop-signal delay, the sessions were divided into four equally sized groups based on the average session SSD: 0-25th SSD percentile, 25-50th SSD percentile, 50-75th SSD percentile, and 75-100th percentile. A separate model was fitted for each trial side and SSD percentile group, with SSRT as dependent variable and RTgo as independent variable. Variables were standardized to obtain comparable standardized beta coefficients.

If it were true that going and stopping share motor dynamics, it is reasonable to expect that sessions where animals needed more time to respond on go trials would also have longer stopping times. For that reason, we tested the hypothesis of shared motor dynamics with linear regression for each trial side separately (*fitlm*, with bisquare weighting function), with RespTgo as independent variable and SSRT as dependent variable. Variables were standardized, so we could average the standardized beta coefficients among animals with significant regressions.

When animals are attentive they are likely to respond quickly to the initial signal and perform well as they are paying attention to the location of the stimulus. As such, a possible effect of attention was tested by comparing SSRTs between sessions with high average RTgo and low response accuracy on go trials (low attention) and sessions with low average RTgo and high response accuracy on go trials (high attention). Sessions belonged to the low attention group if the average RTgo was higher than the median across all sessions (median RTgo_left_ = 254.0 ms; median RTgo_right_ = 234.9 ms) and the average response accuracy on go trials was lower than the median response accuracy across all sessions (median Ago_left_ = 86.0%; median Ago_right_ =87.6%). Sessions belonged to the high attention group if the average RTgo was lower than the median across all sessions and response accuracy on go trials was higher than the median response accuracy across all sessions. A t-test for random samples was used for each trial side separately to statistically test whether attention (low vs. high) had a significant effect on SSRT.

## Results

In the following sections, we will address three main questions. 1) Are single-session SSRTs statistically meaningful as compared to the across-session SSRT? 2) Which factors play a role in the reliability of single-session SSRT estimates? 3) Which factors contribute to varying SSRTs?

## Trends in single-session SSRT estimates

From visual inspection (Figure 5) it is clear that the single-session SSRTs varied over sessions (each session is a different testing day). We first asked whether the SSRTs followed a temporal trend that could account for this variability, e.g., if SSRTs decrease over time as the animals were in the task for longer (but note that the animals were fully trained before any of the data shown here). As single-session SSRT estimates seemed far from constant, we investigated whether SSRT estimates followed a trend as the number of acquired sessions progressed. Polynomial fitting revealed only non-zero constants, linear and quadratic trends, as additional polynomial orders were not statistically improving the fit (Figure 6). For some animals (blue, orange, brown) SSRTs followed a negative quadratic trend (n-shaped), while for other animals (green, brown) the SSRTs followed a positive quadratic trend (u-shaped). For one animal (brown) the sign of the quadratic fit was opposite between left and right SSRTs, while for another animal (green) the sign was similar. Left SSRTs were constant for one animal (yellow), and right SSRTs were constant for two animals (black and blue). After all, SSRT trends were not comparable between animals and none of them had similar kinds of polynomial fits for left and right SSRTs.

**Figure 5.**
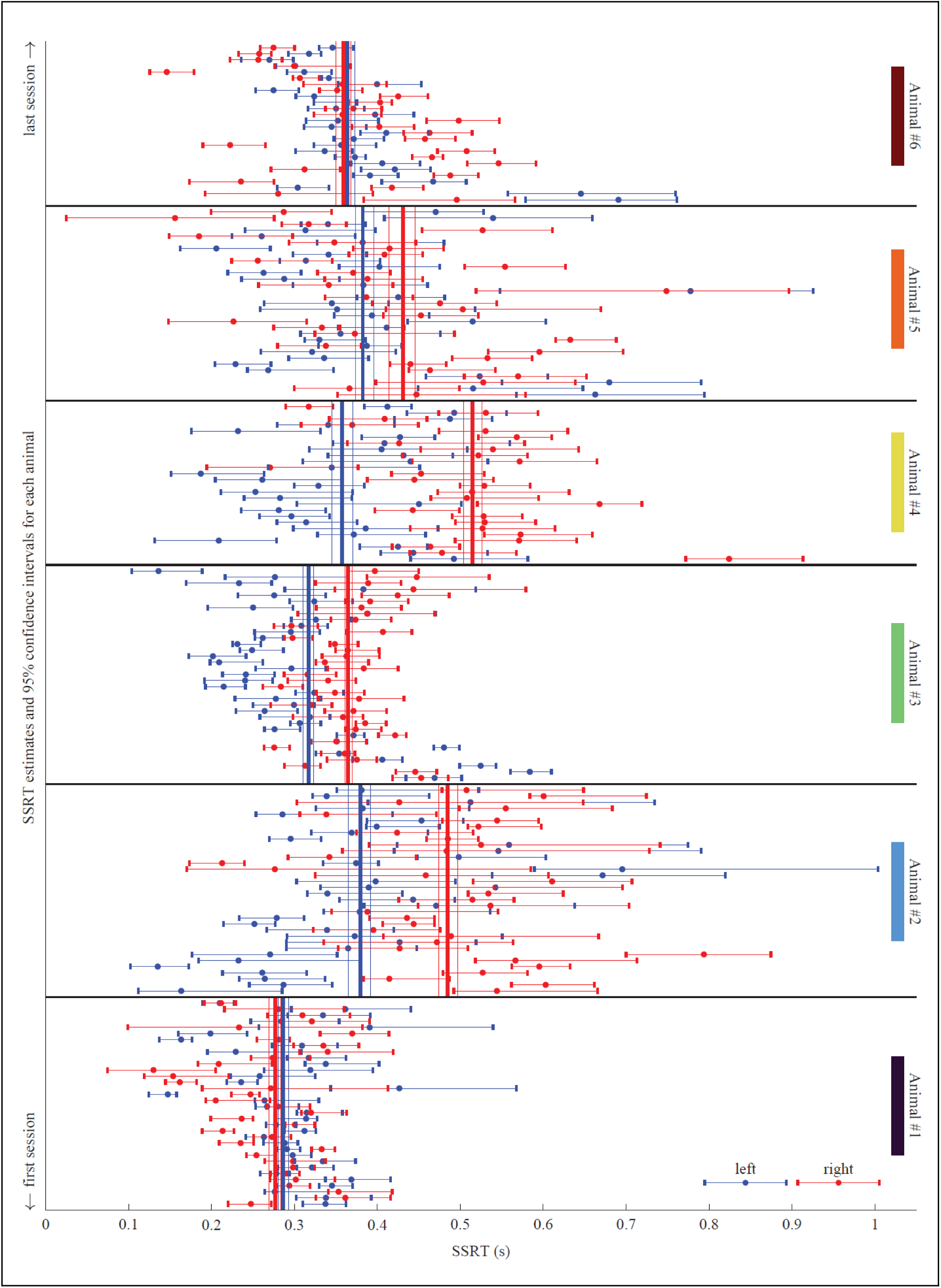
Single-session stop-signal reaction time (SSRT) estimates for each animal, with corresponding 95% confidence intervals. Blue-colored dots represent SSRTs belonging to left stop trials, while red-colored dots represent SSRTs belonging to right stop trials. Vertical blue and red thick lines are across-session SSRT estimates (in case all sessions within an animal would have belonged to one big session) with 95% confidence intervals indicated with thinner vertical lines. Each animal is labeled with a color that matches color-use in other figures.

**Figure 6.**
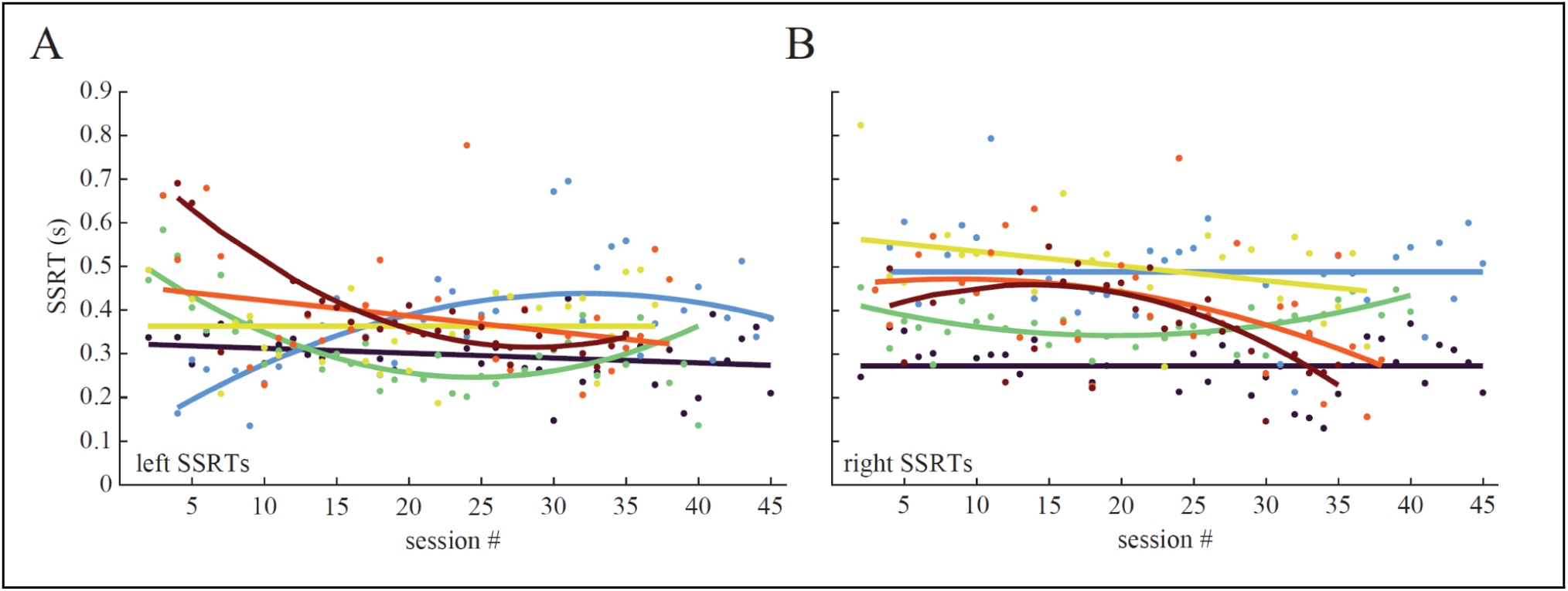
Single-session stop-signal reaction time (SSRT) estimates show different trends over time for different animals and stop-signal sides (**A**, left SSRTs; **B**, right SSRTs). Each dot represents a single session from one animal, where different colors indicate different animals. Trendlines, illustrated with thick colored lines, are polynomials with either a non-zero constant, linear or quadratic fit. For left SSRTs, data from animal #4 (yellow) was best explained with a constant SSRT, while for right SSRTs, data from animal #1 (blue) and #2 (black) were best explained with a constant SSRT. All other groups of SSRTs were better explained by either a linear decline, or negative and positive quadratic polynomials. Note that the polynomial fits are not all spanning the same amount of sessions, because the amount of sessions was not identical between animals.

## Statistical meaningfulness of single-session SSRT estimates

Visual inspection of Figure 5 suggests that individual session SSRTs might be significantly different from the across-session SSRT. To statistically address this, we checked whether confidence intervals of single-session SSRTs overlapped with the across-session SSRT. Confidence intervals of single-session SSRTs overlapped with the across-session SSRT in 47.2% (95% CI [42.9 51.5]) and 43.1% (95% CI [35.4 50.8]) of the sessions for left and right, respectively (see unfiltered SSRT proportions in Figure 7 for individual animal proportions). This means that more than half of sessions had a significantly different SSRT than the across-session SSRT. It is possible that the results described above were simply due to a trend over time, for example if SSRTs decrease over time as animals gained expertise with the task (but note that the animals were fully trained before we started collecting data). Therefore, we applied detrending to the SSRT data with the polynomial fits we showed before (Figure 6) so we were able to correct for trends in SSRTs. Detrending the SSRTs numerically increased the overlap between confidence intervals of single-session SSRTs with the across-session SSRT to 55.2% (95% CI [48.7 61.8]) and 51.5% (95% CI [46.2 56.8]) for left and right, respectively (see detrended SSRT proportions in Figure 7 for individual animal proportions). This and following comparisons between unfiltered and detrended SSRT proportions were not statistically tested because of low sample size (N = 6), which made tests heavily underpowered.

**Figure 7.**
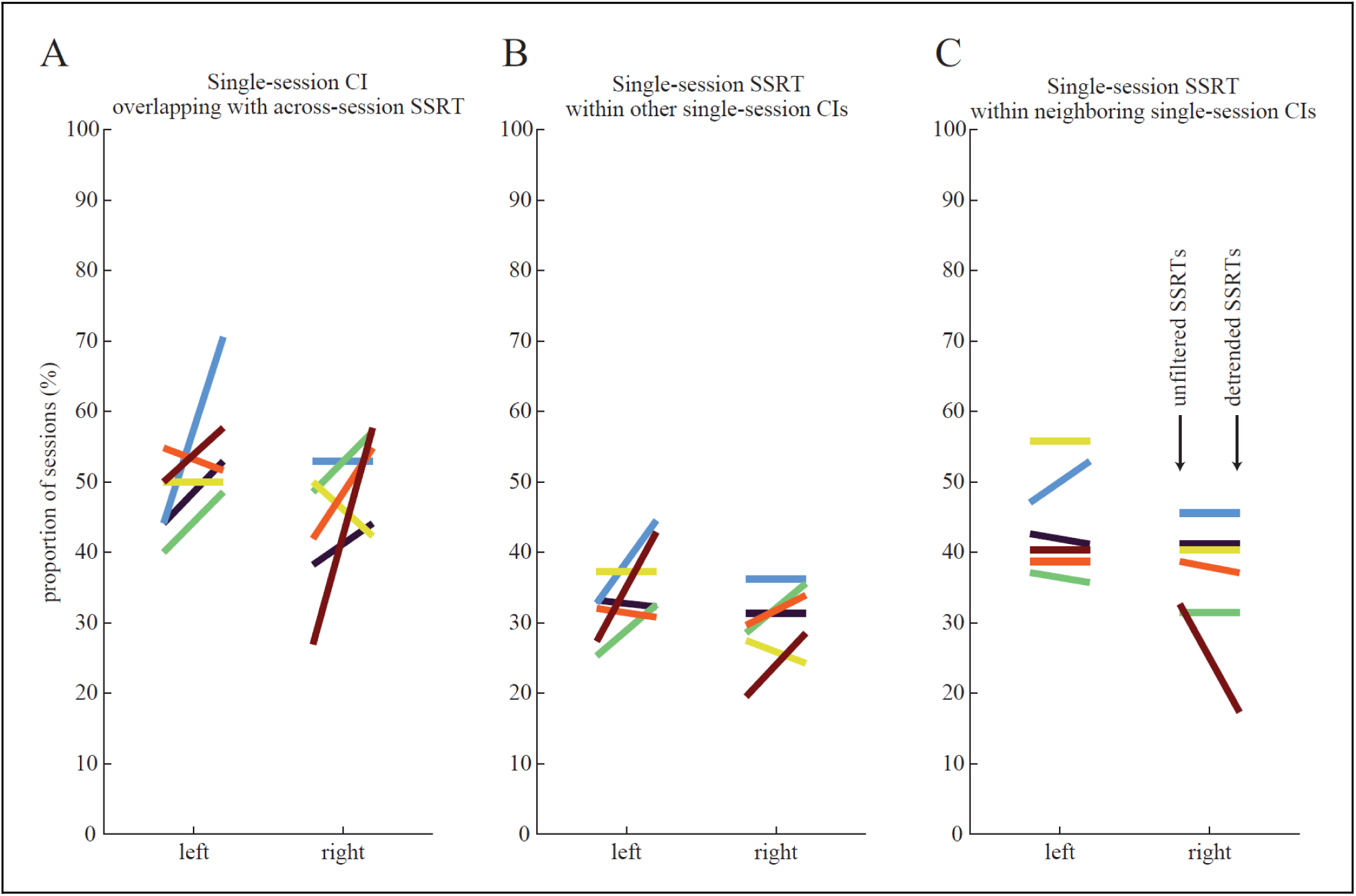
Proportion of sessions meeting three different criteria. **A**, Proportion of sessions where the single-session confidence interval (CI) is overlapping with the across-session stop-signal reaction time (SSRT). **B**, Proportion of sessions where the single-session SSRT is positioned within all other single-session CIs. **C**, Proportion of sessions where the single-session SSRT is positioned within neighboring single-session CIs (see Methods). Different colors represent different animals, and the stop-signal side is separated in two different columns (left/right). Within each left and right separation, two proportions are connected with a colored line. The left proportion is computed without any detrending (unfiltered SSRTs), while the right proportion is computed after detrending single-session SSRT estimates (detrended SSRTs).

To find out how (dis)similar single-session SSRTs were, we computed the average proportion of sessions where the single-session SSRT was positioned within the 95% confidence interval of other single-session SSRT estimates. For unfiltered SSRT data, only 31.3% (95% CI [34.8 27.8]) of left and 28.8% (95% CI [24.4 33.2]) of right single-session SSRTs were statistically similar to other single-session SSRTs, indicating that the majority of single-session SSRTs were not coming from the same distribution. Detrending the SSRTs numerically increased the overlap of neighboring sessions to 36.7% (95% CI [32.0 41.4]) and 31.7% (95% CI [28.0 35.3]) for left and right, respectively.

As one might expect proximate or neighboring sessions to have comparable SSRTs when SSRTs are slowly changing over time, the latter analysis was also applied for neighboring sessions only to investigate the degree to which neighboring sessions are (dis)similar. With the unfiltered SSRT data, this resulted in an increased fraction of SSRTs positioned within single-session confidence intervals, i.e. 43.6% (95% CI [40.1 47.1]) of left and 38.3% (95% CI [33.9 42.7]) of right single-session SSRT estimates were statistically similar to other single-session SSRTs. Although this increase indicates that neighboring sessions had more comparable SSRT estimates than distant sessions, it still shows that more than half of the single-session SSRTs were statistically significantly distinct from the surrounding SSRTs. Detrending the SSRTs numerically increased this to 44.1% (95% CI [37.6 50.7]) for right SSRTs, and numerically decreased to 35.5% (95% CI [27.4 43.6]) for left SSRTs.

## Reliability of single-session SSRT estimates

Estimating the non-observable SSRT from smaller than ideally-sized sessions came with less reliable estimations. Separately computing the left and right SSRT reduced sample size even more for each SSRT estimation. Therefore, we explored which factors relate to the degree of reliability of the SSRT estimate, by using the width of the 95% confidence interval as a dependent variable in a set of linear regression models for each animal and trial side separately (Figure 8). Across both trial sides, three out of five variables showed a significant group effect: variability of go trial response times (mean β*_left_ = .47, *p* = .031; mean β*_right_ = .53, *p* = .031), skewness of go trial response time distribution (mean β*_left_ = -.35, *p* = .031; mean β*_right_ = -.33, *p* = .031) and stop accuracy (mean β*_left_ = -.33, *p* = .031; mean β*_right_ = -.40, *p* = .031). All animals had an individual significant effect of variability of go trial response times for left and right trials, while go trial response time distribution skewness was not significant in three models: 1^st^ animal left (dark blue), 1^st^ animal right (dark blue) and 5^th^ animal right (orange). Stop accuracy did not have an individual significant effect in one model: 3^rd^ animal right (green). Despite the absence of these four individual significant effects, the signs of standardized beta coefficients were in line with the other animals. For all animals, at least two out of five, but not more than three out of five, variables significantly contributed to the explained variance of SSRT reliability.

**Figure 8.**
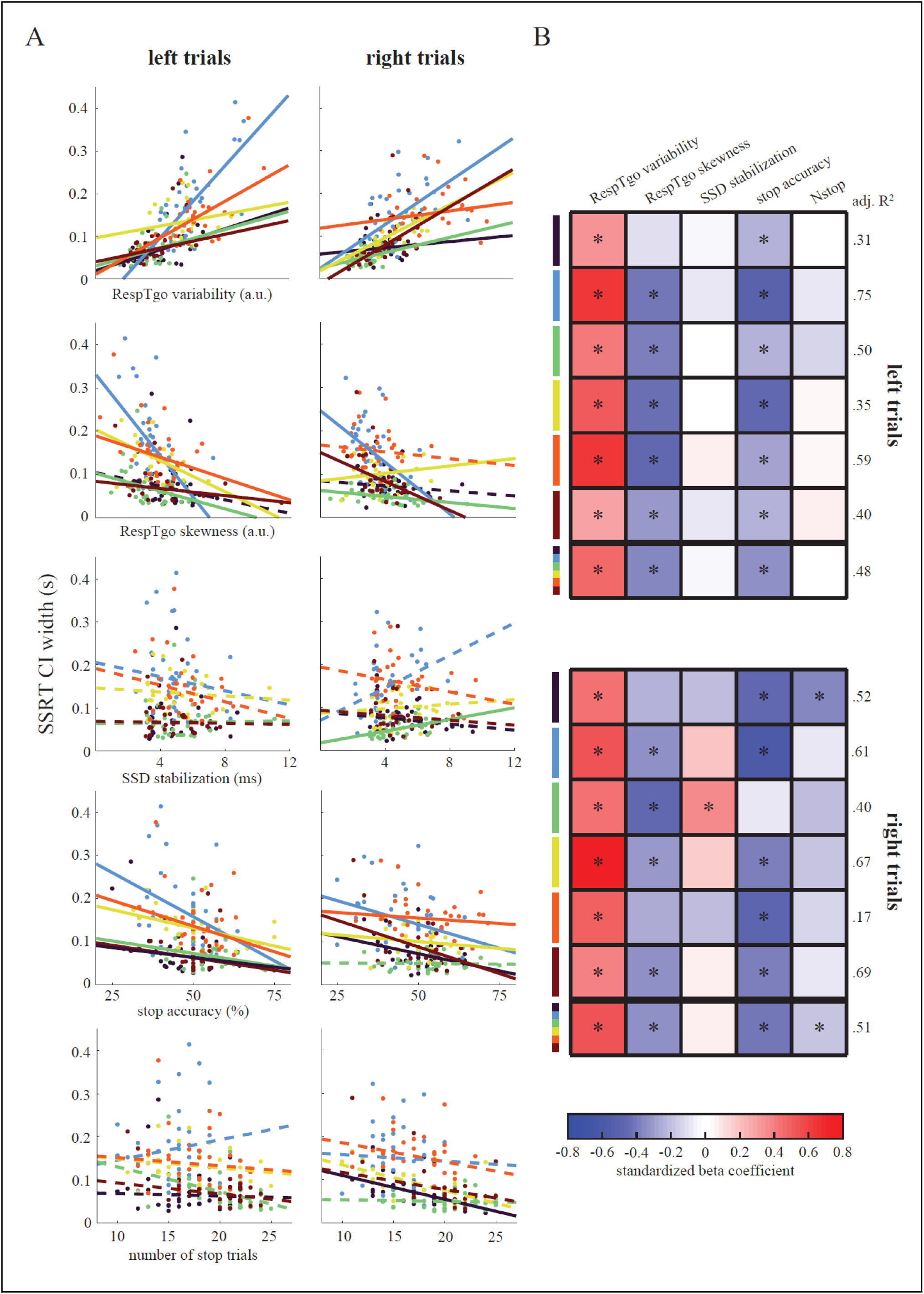
Robust linear regression with stop-signal reaction time (SSRT) 95% confidence interval (CI) width as a proxy for SSRT estimate reliability. **A**, Scatterplots for each independent variable (RespTgo variability, RespTgo skewness, SSD stabilization, stop accuracy, number of stop trials) and SSRT CI width, for left and right trials separately. Each dot represents a session and colors represent different animals. Linear lines are extracted from simplified robust linear regression models (see Methods), and are solid when *p* < .05 in the full robust linear regression model, and dashed in case of non-significance. **B**, Summary heatmaps for left and right trials separately, in which each of first six rows represents an animal, containing the standardized beta coefficients for each independent variable (columns) in the full robust linear regression model. Asterisks indicate *p* < .05 of that predictor variable in robust linear regression, and match with solid lines in panel A. Last row in each heatmap contains average standardized beta coefficients across all six animals, and asterisks indicate *p* < .05 in the Wilcoxon signed-rank test. Colored bars on the left of the heatmap represent different animals and match the color-scheme of panel A and other figures in this chapter. Adjusted R^2^ values from the full model are shown on the right side of the heatmap. RespTgo = response time on go trials; SSD = stop-signal delay.

The Wilcoxon signed-rank test for group significance was significant only when signs of standardized beta coefficients were consistent across all six animals, which resulted in an additional significant group effect for number of stop trials for right trials (mean β*_right_ = -.18, *p* = .031). However, only one animal had individual significance for this variable. On average, roughly half of the SSRT’s variance in reliability was explained by the robust linear regression model (adj. R^2^_left_ = .48; adj. R^2^_right_ = .51). On an individual animal basis, all robust linear regression models were significantly different from a constant model (all *p* < .05) except for the right trial model of the 5^th^ animal (orange; *F_right_*(31, 25) = 2.24, *p* = .082). In line with this, the lowest amount of SSRT reliability variance was explained for the 5^th^ animal (orange; mean adj. R^2^ = .38), while the highest amount of variance was explained for the 2^nd^ animal (light blue; mean adj. R^2^ = .68).

## SSD and nthRespTgo contribution to SSRT estimates

Higher values of nthRespTgo were not necessarily accompanied with higher SSDs, and higher SSDs were not always accompanied with higher values of nthRespTgo (Figure 9). Thus, slower movement speeds were not associated with later stop-signal onsets and vice versa. This is exactly what could have been expected from session-to-session changing SSRTs. When SSRTs would have been similar across sessions, a changing movement speed would have always been perfectly compensated with a changing SSD to obtain a similar SSRT (resulting in dots lined-up on the diagonal across-session SSRT in Figure 9). The average difference across animals between single-session SSRTs and the across-session SSRT was 73.9 ms (95% CI [58.6 89.2]) and 71.3 ms (95% CI [50.4 92.1]) for left and right trials, respectively.

**Figure 9.**
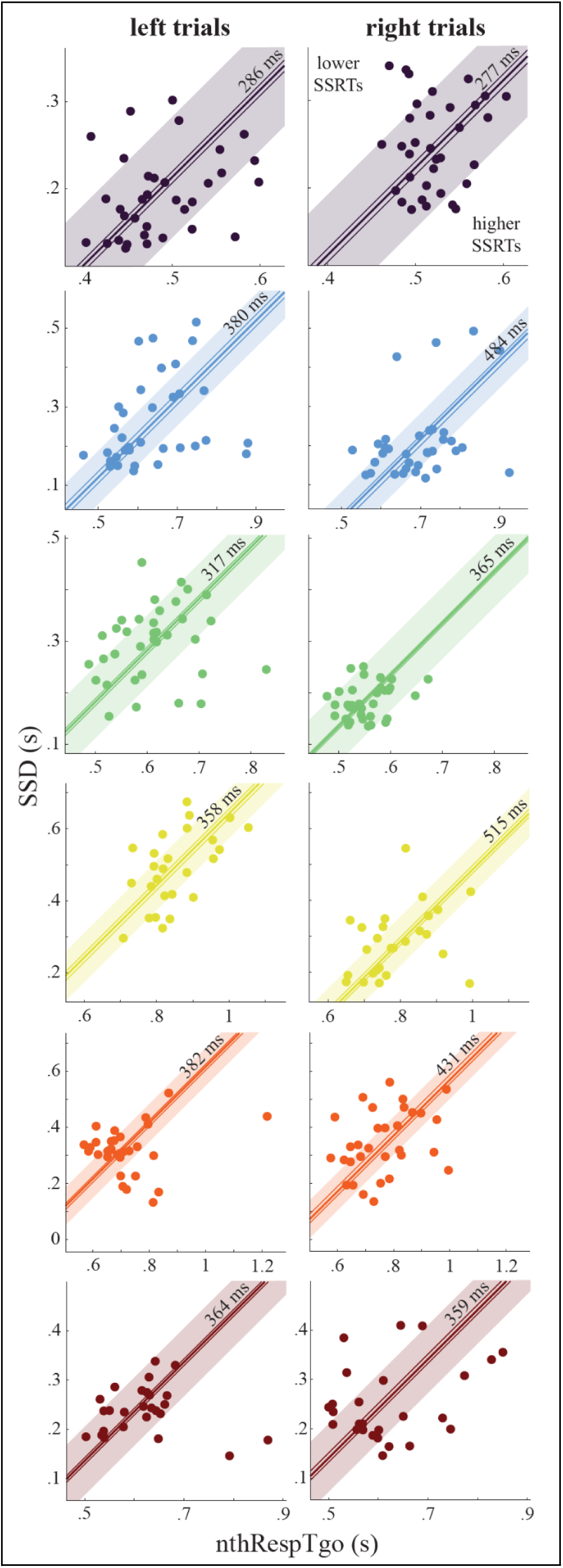
Stop-signal delay (SSD) and nthRespTgo in relation to the across-session stop-signal reaction time (SSRT) estimate. Each animal is indicated with a different color. Each dot is a session and corresponds to a single-session SSRT estimate (SSRT = nthRespTgo - SSD). X- and y-axis limits are identical within each animal, so dot positions and across-session SSRTs between trial sides (left/right) are directly comparable. The diagonal line represents the across-session SSRT estimate (value also printed close to each diagonal line), and includes 95% confidence intervals with thinner diagonal lines. Orthogonal distance from dot to diagonal SSRT line represents deviance from across-session SSRT. As such, all positions on the diagonal line are equal to the across-session SSRT, while dots left of and above the line represent sessions with a lower SSRT than the across-session SSRT, and dots right of and below the line represent sessions with a higher SSRT than the across-session SSRT. Width of the shaded area is equal to 100 ms (50 ms on each side of across-session SSRT). nthRespTgo = n-th response time on go trials matching p(failed stopping).

SSRT was computed by subtracting SSD from nthRespTgo. Since both observable variables had considerable spread across sessions, it was not a surprise that robust linear regression revealed that both variables were significant predictors for SSRT for all animals and trial sides (all *p* < .001). The main goal of fitting the robust linear regression model was to extract standardized beta coefficients, so we could directly compare each variable’s contribution to SSRT. While for some animals absolute standardized beta coefficients were numerically bigger for nthRespTgo than for SSD, for some animals this was the opposite (mean β*_nthRespTgo,left_ = 0.86 (ranging from 0.80 to 0.95), mean β*_SSD,left_ = -0.79 (ranging from -1.15 to -0.50); mean β*_nthRespTgo,right_ = 0.85 (ranging from 0.59 to 0.94), mean β*_SSD,right_ = -0.83 (ranging from -0.92 to -0.72)). A paired-samples t-test indicated that there were no statistically significant differences between absolute standardized beta coefficients of SSD and nthRespTgo, for both left and right trials (t_left_(5) = .87, *p* = .424; t_right_(5) = .17, *p* = .873), meaning SSD and nthRespTgo had statistically comparable contributions to SSRT estimates.

## Possible cognitive and neural mechanisms driving session-to-session changes in SSRT

Because SSRTs changed from session to session, we reflected on which cognitive and neural factors could drive these changing SSRTs. We identified three, in our opinion, most likely contributors: Motivation for quick rewards, shared motor dynamics and attention. We attempted to quantify these cognitive and neural mechanisms with variables from the task, and theoretically addressed other possible driving factors in the discussion which we could not approximate with task variables.

### Motivation for quick rewards

It is possible that sessions with faster SSRTs were those in which animals were more motivated to obtain rewards (e.g., they were thirstier in some sessions). If this were a viable explanation, it would predict that the release time on go trials (RTgo) would strongly correlate with SSRT. Therefore, we regressed RTgo with SSRT in four equally sized SSD quantiles across animals for left and right trials separately (Figure 10A). Almost all quantiles showed a significant positive association between RTgo and SSRT (except for 3rd and 4th quantile of left trials, all other *p* < .05). Across trial sides (left/right), standardized beta coefficients and adjusted R^2^s were smaller for each next SSD quantile (β*_1st_ = 0 .61, β*_2nd_ = 0.58, β*_3rd_ = 0.32, β*_4th_ = 0.16; adj. R^2^_1st_ = 0 .41, adj. R^2^_2nd_ = 0.31, adj. R^2^_3rd_ = .09, adj. R^2^_4th_ = 0.04), suggesting that high motivation for quick rewards (as reflected with low RTgo) was associated with fast stopping, but mostly in sessions where stop-signals were presented relatively early, and not as much when stop-signals were presented later.

**Figure 10.**
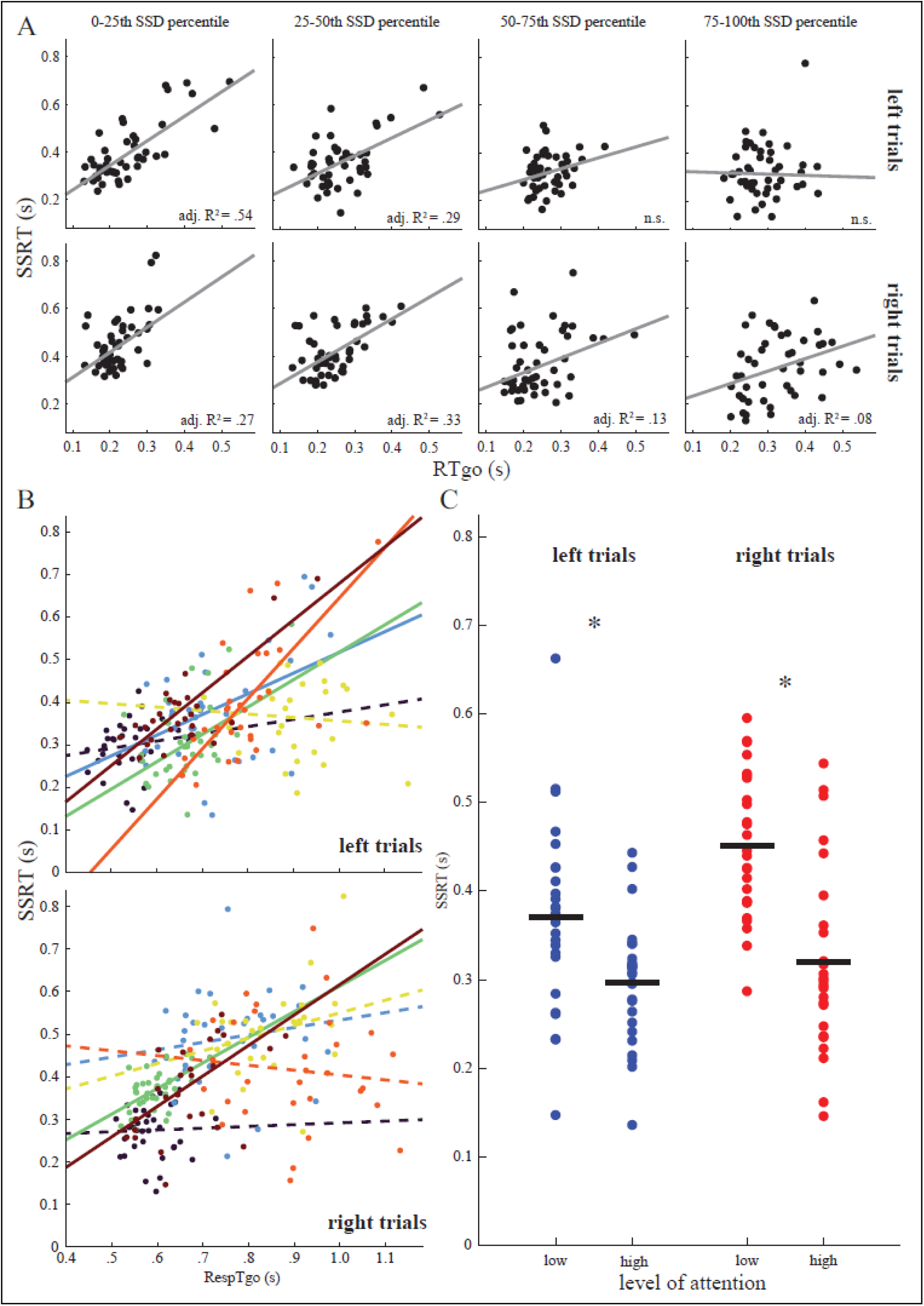
Motivation, motor dynamics and attention contribute to session-to-session changing SSRTs. **A**, Linear regression between RTgo and SSRT for 1st, 2nd, 3rd and 4th percentile group of session SSDs across all six animals (therefore no colored dots). Dots represent different sessions. Left and right trials separated by top and bottom row, respectively. Grey line represents the robust linear fit. Adjusted R^2^ printed in right bottom corner if *p* < .05, otherwise non-significant (n.s.). **B**, Linear regression between RespTgo and SSRT for each animal and trial side separately. Dots represent different sessions. Different animals are indicated with different colors, with the same color scheme as the other figures in this chapter. Dashed lines indicate non-significant regressions, while solid lines indicate significance (*p* < .05). **C**, Across animals, sessions with high attention have significantly lower SSRTs as compared to sessions with low attention. Blue dots are left trials, red dots are right trials. Black horizontal lines indicate the average SSRT of that group. Asterisks indicate significance (*p* < .01). RTgo = release time on go trials; SSRT = stop-signal reaction time; RespTgo = response time on go trials.

### Shared motor dynamics

As elements of motor dynamics between going and stopping could be shared, we assessed whether RespTgo and SSRT were positively associated (Figure 10B). For left trials, 4 out of 6 animals had a significant positive association between RespTgo and SSRT (mean β*_left_ = 0.63, all *p <* .05), while for right trials only 2 out of 6 animals had a significant positive association (mean β*_right_ = 0.59, all *p <* .05). Therefore, general motor dynamics were likely also a contributor to across session SSRT variability.

### Attention

We approximated the level of attention with a combination of two variables: RTgo and response accuracy on go trials (Figure 10C). Sessions in which animals had relatively low attention (high RTgo and low accuracy on go trials) had significantly higher SSRTs (mean SSRT_left_: 371.1 ms; mean SSRT_right_: 451.3 ms), as compared to sessions where animals had relatively high attention (low RTgo and high response accuracy; mean SSRT_left_: 296.6 ms; mean SSRT_right_: 319.7 ms) (*t_left_*(54) = 3.2583, *p* < .01, Cohen’s *d* = 0.87; *t_right_*(50) = 5.0951, *p* < .001, Cohen’s *d* = 1.41). Therefore, sessions with higher levels of attention were also associated with faster stopping speeds.

## Discussion

In this study, we collected many sessions per animal in a rodent version of the stop-signal task, and explored whether single-session SSRT estimates were statistically meaningful as compared to a single SSRT estimate across all sessions. We showed that stopping speeds were changing from session to session and did not show a basic or consistent trend over time across individuals. As single-session SSRT estimates came with higher degrees of estimate uncertainty, we wanted to know which factors were underlying these increasing uncertainties. Higher degrees of single-session SSRT reliability were associated with lower go trial response time variabilities, lower skewnesses of the go trial response time distribution and higher stop accuracies. SSRT variability is equally explained by variability in stop-signal timing and go trial response times, and motivation, shared motor dynamics and attention could partly explain changing stopping speeds. This study suggests that stopping speed is a state that can meaningfully change from time to time, and provides insights for researchers that are interested in collecting multi-session stop-signal task data.

## Stopping speed is a state, not a trait

Across the six animals and within each animal, there was a lot of variability in stopping times. Before we quantified how variable and dissimilar they were to each other, we examined whether consecutive single-session SSRTs were following a consistent logical trend. Trend analysis revealed that SSRTs were not consistently following a pattern, across animals and within-animal across stop-signal sides. Because changing SSRTs were not simply explained by something like an increasing or decreasing trend, we determined how dissimilar they were as compared to the across-session SSRT. We found that more than half of the single-session estimates’ confidence intervals were not coinciding with the across-session SSRT, only a minority of single-session SSRTs were likely to share the same SSRT with other sessions, and neighboring sessions had only slightly more similar stopping times. Trends in SSRTs only marginally explained these measures, as on average it did not even increase the overlap between within-session SSRTs and across-session SSRTs with 10%. These findings suggest that most single-session SSRT estimates are not only numerically, but also statistically different from an across-session, trait-like SSRT. Altogether, this supports the hypothesis that stopping speed is a state-like characteristic that meaningfully changes over time. Previous literature has already shown that within-subject stopping speeds can be different when task designs are different (Gordi et al., 2019; Doekemeijer et al., 2023; Weber et al., 2024), but to our knowledge this is the first report that rigorously demonstrates to what degree and partly why stopping speeds change under identical experimental conditions.

## SSRT reliability correlates with different task parameters

Sessions with higher variability in go trial response times were associated with lower levels of SSRT reliability. This could be explained by the possibility that the tracking procedure of stop-signal delays is less effective in such sessions. If response speeds are varying on the order of 100s of milliseconds across trials, but the stop-signal delay is limited to adaptations of 50 ms after each stop trial, it may be that this negatively impacts the reliability of the SSRT estimate. For example, an SSD of 350 ms in one trial may be very easy in a trial with slow going speed, but disproportionately more difficult if the SSD changes to 400 ms while the rat goes 300 ms faster the next stop trial. This could make the tracking procedure less effective, in turn affecting the reliability of the SSD used for estimating the SSRT.

Increased skewness of the go trial response time distribution is thought to affect SSRT reliability negatively (Verbruggen et al., 2019), as it may reflect slowing behavior over the course of a session in an attempt to outsmart the tracking procedure (which increases the skewness of the go trial response time distribution), even though this does not pay off with an adaptive SSD (Verbruggen et al., 2013). To our surprise, we found that sessions with higher levels of skewness had more reliable SSRTs. Our speculation is that the estimate of skewness may have been negatively impacted by low numbers of trials contained in the response time distribution, possibly by allowing relatively high response times to have disproportionate effects on skewness, as this estimate is susceptible to outliers. However, future studies may shed light on whether increased skewness always has positive effects on SSRT reliability in multi-session stop-signal tasks, or whether this relationship is mediated by another unknown factor.

As expected, lower accuracies on stop trials negatively impacted SSRT reliability. The stop-signal task is designed to get as close to 50% accuracy on stop trials as possible to increase the probability of obtaining reliable SSRTs, even when using the integration method (Verbruggen et al., 2019). Although not explicitly tested here, it seems that stop accuracies at the higher end of the spectrum were not necessarily improving the SSRT reliability substantially, which may reflect a ceiling effect. Altogether, across-animal analysis showed that higher go trial response time variabilities, lower levels of skewness in the go trial response time distribution and lower stop trial accuracies negatively impact the reliability of SSRT estimates in our multi-session stop-signal task.

## Stop-signal timing and going speed equally contribute to SSRTs

Before we addressed cognitive and neural mechanisms that could possibly drive changing SSRTs, we investigated whether stop-signal timing and going speed were equally contributing to SSRT estimations or not, as they are both variables that make up the SSRT by subtracting the session SSD from the n-th go trial response time that matches 50% performance in that session. Although one might intuitively think that they are contributing equally by definition due to simple subtraction, this is not necessarily the case because one variable may vary substantially more from session to session than the other. For some animals variance in one variable explained more SSRT variance than the other variable, but for other animals this was the opposite. However, across animals, we statistically and visually demonstrated that variance in SSD and nthRespTgo equally contributed to varying SSRTs. Sessions in which animals were going slower were not always accompanied with equally delayed stop instructions, meaning they were slower stoppers in those sessions. Similarly, sessions in which animals were going faster were not always accompanied with equally earlier stop instructions, meaning they were faster stoppers in those sessions. This again supports the notion that SSRTs are not constant, but change from session to session.

## Stopping speed may vary with changing motivation, going speed and attention

When animals were highly motivated to get a reward quickly (as indicated with low release times on go trials), for example because they were thirsty, they also had faster stopping speeds. However, this relationship was observed for sessions with early stop-signal presentations (low SSDs), and to a lesser degree in sessions where stop-signals were presented relatively late. This implies that rats were mostly stopping faster during highly motivational states in sessions where stop-signals were presented fairly early, but not when presented late. When comparing this to the traffic light example, this would mean that you are well capable of stopping quickly when you are in a rush to reach your destination if the traffic light turns red quite distant from the intersection. But, in comparable rushing circumstances, you turn out to be a slower stopper when it turns red when you are closer to the intersection. Why would this happen? And specifically, why would proximity to the intersection impact stopping speed? We speculate that highly motivated individuals possess some degree of impatience that affects their sensitivity to late-presented stop-signals. Some time after starting to go they may just decide (consciously or unconsciously) to not process new incoming visual stimuli, as they focus solely on getting a reward as quickly as possible. This cognitive tunneling may specifically slow down stopping speed in sessions with late-presented stop-signals, due to increasing cognitive load the later a stop-signal is added to the already focused state of mind. Another thought could be that sessions in which rats release early from the central port have more likelihood of stop-signals being presented late in the go-process, increasing the likelihood of failing at stopping in time. However, in this situation the SSD would decrease in turn to compensate for poor performance on stop trials. Moreover, in sessions where the average SSD and RTgo are relatively low, the positive relation between motivation and stopping speed is strongest.

For some animals, faster going speeds were associated with faster stopping speeds. We want to stress that this effect was not present for every animal, implying that some animals support the hypothesis of shared motor dynamics between going and stopping, while other animals do not support this hypothesis. However, no animal’s data supported the hypothesis that going and stopping speeds were negatively correlated. In chapter 3 of this dissertation (Figure 2C), we also investigated this relationship and included more animals and more sessions. Here we did observe a significant positive correlation between going speed and stopping speed across animals and trial sides (left/right). Future multi-session stop-signal task studies have to elucidate whether this relationship is replicable.

Sessions in which animals were highly attentive, as indicated by fast release times and high response accuracies on go trials, were accompanied with faster stopping times as compared to sessions where animals displayed signs of lower attentive states, as approximated with slower go trial release times and response accuracies. As with many cognitive tasks, adequate attention to a relevant stimulus is key for trial outcome success. Although *attention* could have many definitions, it is generally associated with better performance in combination with faster reaction times (Carlson et al., 1983; Prinzmetal et al., 2005). In our task, rats had to divide their attention to both the left and right visual field to be able to quickly release from the central port after go-signal onset and respond correctly. Response accuracy would only reach high levels when release from the central port (marking RTgo) was not just triggered by non-spatially noticing the go-signal, but by being attentive to the spatial location of the go-signal. Our data suggest that rats are also more attentive to a potential stop-signal when they are (spatially) attentive to the go-signal. Translated to a real-world human example this would mean that red traffic lights can be presented later if you are anticipating the emergence of the stop-signal, and still be able to stop in time because your stopping speed increases with increasing attention to the red traffic light.

## Other possible explanations for changing stopping speeds

One could argue that the level of training or developmental age may play a role in changing stopping speeds. However, trend analysis of SSRTs demonstrated that stopping times did not consistently decrease with time across animals (Figure 6). Also, our animals reached a certain training level before they underwent surgery for electrode implantation. After that, the behavioral data were collected, so there is no data available with which we can check whether fairly young rats with not fully developed frontal cortices are slower at stopping than fully-developed rats.

Another rationale would be that a speed-accuracy trade-off affected stopping speeds. However, when rats decide to slow down to increase the likelihood of responding correctly on stop trials (although it results in longer waiting times for a reward), our adaptive procedure automatically tunes the stop-signal delay to a level where they will have equally difficult times with stopping as in sessions where they are moving faster. Successful stopping is followed by a later onset of the stop-signal, meaning it becomes more difficult to stop in time, while failed stopping is followed by an earlier onset of the stop-signal, resulting in a higher probability of stopping in time. On the contrary, when rats decide to increase their speed to shorten the waiting time of receiving a reward (while diminishing the probability of getting one), this will also tune the stop-signal delay to equally difficult levels as a result of how well the rat performs on stop trials. For go trials, adjusting the going speed is rewarding in our stop-signal task, as slowing down simply decreases the likelihood of making erroneous responses (unless they are inattentive). But for stop trials, adjusting going speed is not rewarding due to the adaptive nature of the stop-signal delay based on stopping performance. In fact, we cannot think of a reason why favoring speed or accuracy would impact stopping speed in particular. One could argue that going faster makes stopping more difficult, as there is a cognitive emphasis on going rather than stopping, but we already learned that going faster is not associated with stopping slower.

## Limitations and future directions

The task we used was not optimized for extracting which factors contribute to changing SSRTs and SSRT reliability, as it was designed for obtaining as reliable SSRT estimates as possible. Nevertheless, with our analyses that were mostly correlative of nature, we attempted to shed some light on the statistical significance and reliability of single-session SSRTs. This study could be a starting point for researchers that are interested in using multi-session stop-signal task designs to gain more insights about reactive stopping.

However, the sample size of six animals, while providing initial insights, limited statistical power and the ability to generalize findings. Future studies should consider bigger sample sizes to validate these preliminary findings. Moreover, the relationships between factors that are associated with SSRT reliability remain complex and somewhat contradictory to previous literature. While we speculated that low trial numbers might have disproportionately affected skewness, studies with larger trial numbers and experimentally controlled conditions are needed to clarify these relationships. Another limitation lies in the exploration of cognitive mechanisms underlying SSRT variability. While we identified potential contributions from motivation, shared motor dynamics, and attention, these factors were not directly manipulated or measured in a way that allowed us to make causal inferences. Future studies should incorporate more direct assessments of these cognitive states, perhaps through manipulations, to better understand their influence on stopping speed.

In conclusion, while our findings demonstrate that stopping speeds are not fixed within-animal, future research with larger sample sizes, more controlled experimental conditions, and direct measures of cognitive states will be essential for advancing our understanding of reactive stopping in the context of multi-session approaches. Despite these limitations, we can solidly argue that in a multi-session stop-signal task design, researchers should use *single-session* SSRT estimates instead of an *across-session* SSRT estimate, as within-animal stopping speed substantially and meaningfully changes across sessions. This could turn out to be very useful when the stop-signal task is combined with neural recordings such as local field potentials, because single-session SSRT estimates allow one to time-lock to the SSRT of that session specifically to extract stop-related neural signatures in a more time-precise manner.

## Acknowledgements

J. ter Horst was funded by the Donders TopTalent PhD scholarship. We would like to thank Sjef van Hulten and Arthur de França for technical support and Marie-Anna Sedlinská, Kasif Işik and Mauricio Diaz-Ortiz Jr. for help with training animals and data collection.

## References

Carlson JS, Jensen CM, Widaman KF (1983) Reaction time, intelligence, and attention. Intelligence 7:329–344. doi:10.1016/0160-2896(83)90008-9

Curley LB, Newman E, Thompson WK, Brown TT, Hagler DJ, Akshoomoff N, Reuter C, Dale AM, Jernigan TL (2018) Cortical morphology of the pars opercularis and its relationship to motor-inhibitory performance in a longitudinal, developing cohort. Brain Struct Funct 223:211–220. doi:10.1007/s00429-017-1480-5

Doekemeijer RA, Dewulf A, Verbruggen F, Boehler CN (2023) Proactively Adjusting Stopping: Response Inhibition is Faster when Stopping Occurs Frequently. J Cogn 6:22. doi:10.5334/joc.264

Gordi VM, Drueke B, Gauggel S, Antons S, Loevenich R, Mols P, Boecker M (2019) Stopping Speed in the Stop-Change Task: Experimental Design Matters! Front Psychol 10:279. doi:10.3389/fpsyg.2019.00279

Logan GD, Cowan WB (1984a) On the ability to inhibit thought and action: A theory of an act of control. Psychol Rev 91:295–327. doi:10.1037/0033-295X.91.3.295

Madsen KS, Johansen LB, Thompson WK, Siebner HR, Jernigan TL, Baaré WFC (2020) Maturational trajectories of white matter microstructure underlying the right presupplementary motor area reflect individual improvements in motor response cancellation in children and adolescents. Neuroimage 220:117105. doi:10.1016/j.neuroimage.2020.117105

Prinzmetal W, McCool C, Park S (2005) Attention: reaction time and accuracy reveal different mechanisms. J Exp Psychol Gen 134:73–92. doi:10.1037/0096-3445.134.1.73

Thunberg C, Wiker T, Bundt C, Huster RJ (2024) On the (un)reliability of common behavioral and electrophysiological measures from the stop signal task: Measures of inhibition lack stability over time. Cortex 175:81–105. doi:10.1016/j.cortex.2024.02.008

Verbruggen F, Chambers CD, Logan GD (2013) Fictitious inhibitory differences: how skewness and slowing distort the estimation of stopping latencies. Psychol Sci 24:352–362. doi:10.1177/0956797612457390

Verbruggen F, Logan GD (2009) Models of response inhibition in the stop-signal and stop-change paradigms. Neurosci Biobehav Rev 33:647–661. doi:10.1016/j.neubiorev.2008.08.014

Verbruggen F et al. (2019) A consensus guide to capturing the ability to inhibit actions and impulsive behaviors in the stop-signal task. eLife 8:1–26. doi:10.7554/eLife.46323

Weber S, Salomoni SE, St George RJ, Hinder MR (2024) Stopping speed in response to auditory and visual stop signals depends on go signal modality. J Cogn Neurosci 36:1395–1411. doi:10.1162/jocn_a_02171

Williams BR, Ponesse JS, Schachar RJ, Logan GD, Tannock R (1999) Development of inhibitory control across the life span. Dev Psychol 35:205–213. doi:10.1037//0012-1649.35.1.205

